# A novel nonsense mutation c.424G>T (p. G142X) in the first exon of XLas leading to osteopetrosis

**DOI:** 10.1101/507111

**Authors:** Xiang Chen, Yang Meng, Ying Xie, Shan Wan, Li Li, Jie Zhang, Bo Su, Xijie Yu

**Affiliations:** Laboratory of Endocrinology and Metabolism, Department of Endocrinology, State Key Laboratory of Biotherapy, West China Hospital, Sichuan University, Chengdu, 610041, China; Laboratory of Pathology, West China Hospital, Sichuan University, Chengdu, 610041, China; Core Facility of West China Hospital, Sichuan University, Chengdu, 610041, China

**Keywords:** osteopetrosis, hypophosphatemia, XLas, parathyroid hormone, pseudohypoparathyroidism

## Abstract

GNAS is one of the most complex gene loci in the human genome and encodes multiple gene products. XLas, the extra-large isoform of alpha-subunit of the stimulatory guanine nucleotide-binding protein (Gas), is paternally inherited. Although XLas can mimic the action of Gas, its significance remains largely unknown in humans. Here we report a patient presented with increased bone mass, hypophosphatemia, and elevated parathyroid hormone levels. His serum calcium was in the lower limit of normal range. DEXA scan revealed progressive increase in the bone density of this patient. Whole exome sequencing of this subject found a novel nonsense mutation c.424G>T (p. G142X) in the first exon of XLas, which was inherited from his father and transmitted to his daughter. This mutation was predicted to exclusively influence the expression of XLas, while may have no significant effects on other gene products of this locus. SaOS2 cells transfected with mutant XLas failed to generate cAMP under parathyroid hormone stimulation, indicating skeletal resistance to this hormone. This subject showed higher circulating SOST, DKK1 and OPG levels, while lower RANKL levels and RANKL/OPG ratio, leading to reduced bone resorption. It is speculated that this patient may belong to a very rare type of pseudohypoparathyroidism with selective skeletal resistance but normal renal tubular response to parathyroid hormone. Our findings indicate that XLas plays a critical role in bone metabolism and GNAS locus should be considered as a candidate gene for high bone mass.

**Author summary:** GNAS has been regarded as one of the most complex gene loci and encodes multiple transcripts, including Gsα, XLαs, NESP55 and A/B transcripts. These isoforms share the same 2-13 exons with alternative first exons. Previously reported mutations often disrupt multiple protein-coding transcripts in addition to that encoding Gsα, making it difficult to distinguish the contributions of each transcript to disease phenotypes. Here we first report a novel nonsense mutation c.424G>T (p. G142X) in the first exon of XLas in a subject presenting with high bone mass, unclosed cranial suture, and persistent hypophosphatemia, and elevated parathyroid hormone (PTH) levels. This is the first report of a mutation located in the first exon of XLas in humans, which was predicted to exclusively influence the expression of XLas, while may have no significant effects on other gene products of this locus. SaOS2 cells transfected with mutant XLas failed to generate cAMP under PTH stimulation, indicating skeletal resistance to this hormone. Our study suggests that XLas has an important physiological role in humans, and is involved in skeletal PTH/cAMP pathway. Our findings also indicate GNAS locus should be considered as a candidate gene for high bone mass.

**Funding:** The National Natural Science Foundation of China.

**Declaration of Interests:** The authors declare no competing interests.

## Introduction

GNAS has been regarded as one of the most complex gene loci in the human genome and undergoes tissue-specific imprinting [1]. Epigenetic event contributes to tissue specific imprinting of Gsα, which leads to phenotypic variability in GNAS mutations. Heterozygous, maternally inherited inactivating mutations in one of the 13 GNAS exons encoding Gsα lead to pseudohypoparathyroidism type 1a (PHP1a). However, paternal inheritance of the same mutations causes pseudopseudohypoparathyroidism (PPHP), which manifests as Albright’s hereditary osteodystrophy (AHO) without hormonal resistance [2].

The GNAS complex locus encodes multiple transcripts, including Gsα, XLαs, NESP55 and A/B transcripts [3]. These isoforms share the common exons 2–13 with four alternative promoters and first exons [4]. Gsα (NM_000516.5) couples seven transmembrane receptors to adenylyl cyclase and is involved in intracellular cAMP generation [5]. In most tissues, Gsα is biallelically expressed. However, only maternal allele of Gsa is expressed in renal proximal tubules. XLαs (NM_080425.3), the extra-large Gsα isoform, is paternally expressed [6]. XLαs shows a more restricted tissue distribution than Gsα, and is primarily expressed in neuroendocrine tissues. Although XLas can mimic the actions of Gsα to stimulate adenylyl cyclase to generate cAMP, obviously different cellular actions have been found between XLas and GSa, possibly due to their different activation-induced trafficking [6–8]. Presently it is unclear whether XLαs has an important physiological function in humans. Here we first report a subject with high bone mass (HBM) carrying a novel mutation in the first exon of XLαs, which indicats that XLas may play a role in bone metabolism.

## Materials and Methods

### Subjects

This study was approved by the Institutional Review Boards of West China Hospital, Sichuan University. All subjects were given written, informed consent before participating in the study. The proband was a 44-year-old man, first seen at the outpatient unit of West China Hospital in May 2014, with complaints of back pain for 12 years, aggravated with double hip pain for 2 years. The patient’s height was 170 cm, and body weight was 62kg. Laboratory evaluation revealed elevated PTH, low serum phosphorous and 25-hydroxyvitamin D levels. His serum calcium was at the lower limit of the normal range. Serum levels of bone specific alkaline phosphatase (BALP), N-mid osteocalcin (OCN) and type 1 collagen cross-linked C-telopeptide (CTX) were increased. He had normal thyroid stimulating hormone (TSH) and free T4 (FT4) levels. His mean bone mineral density, T and Z scores of lumbar vertebrae, L1–L4, were 1.936 (gm/cm^2^), 7.1 and 7.6, respectively. His mean bone mineral density, T and Z scores of femoral neck were 1.406 (gm/cm^2^), 3.3 and 3.8, respectively (table 1). This patient had no history of fluorosis and hepatitis C infection.

**Table 1.**
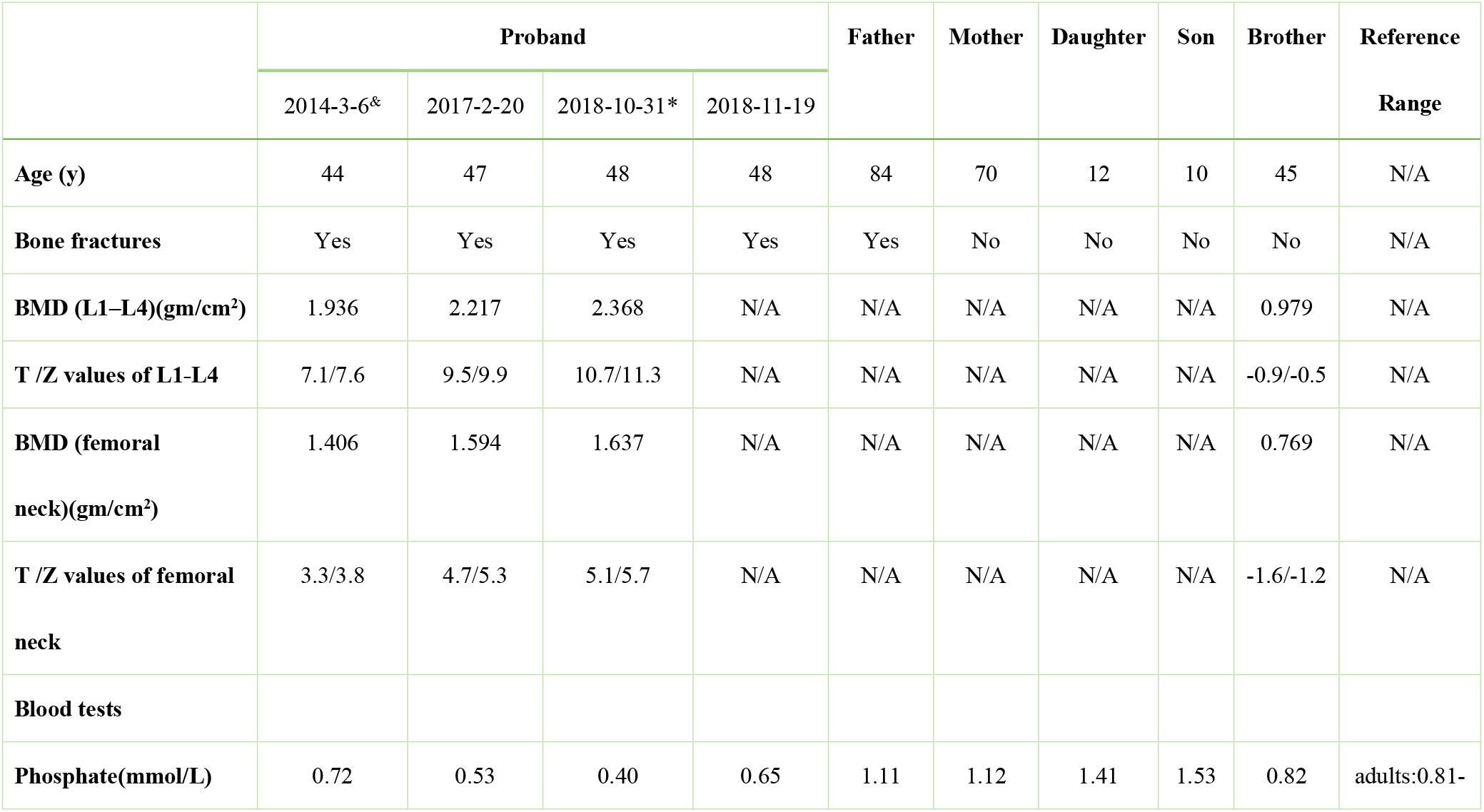
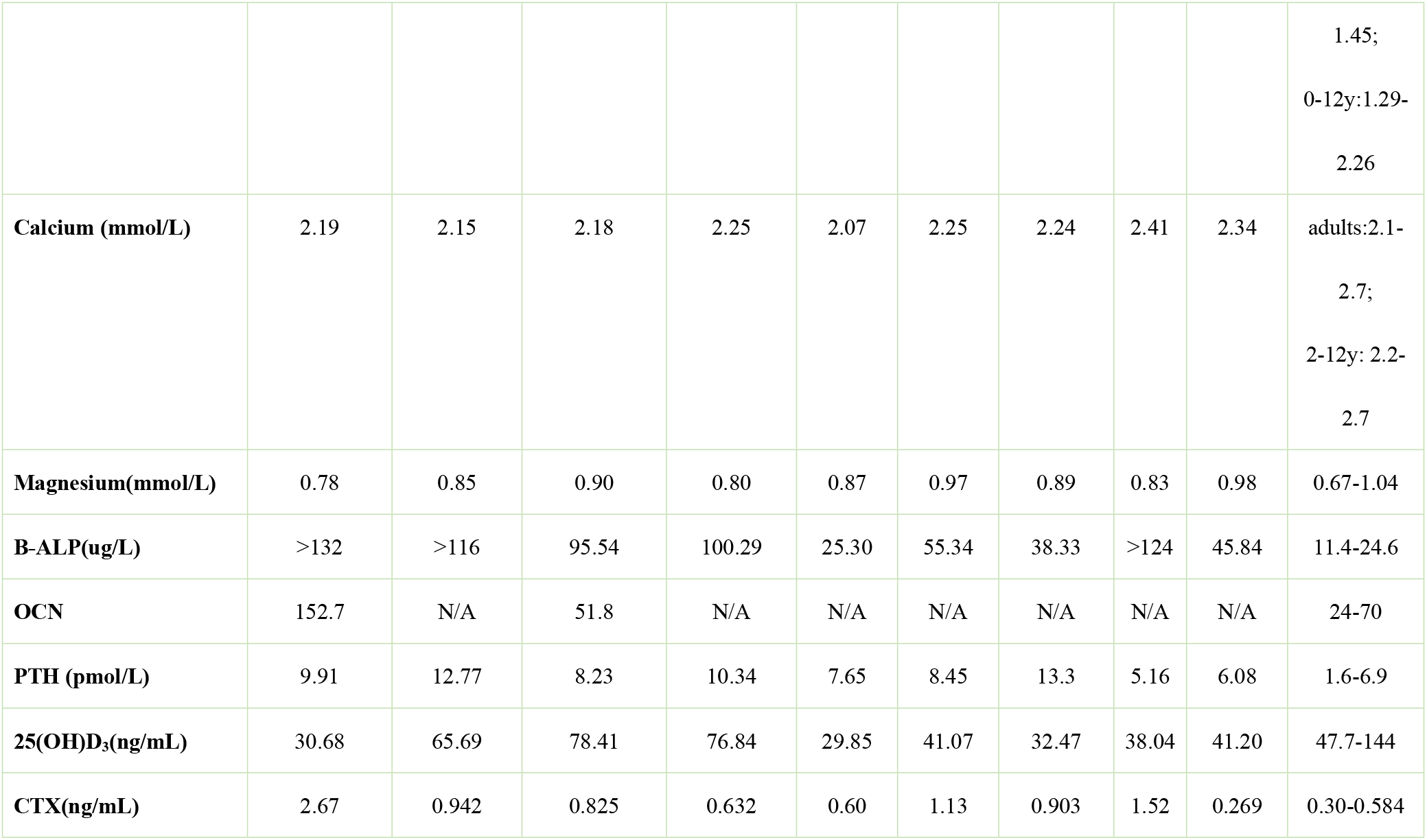
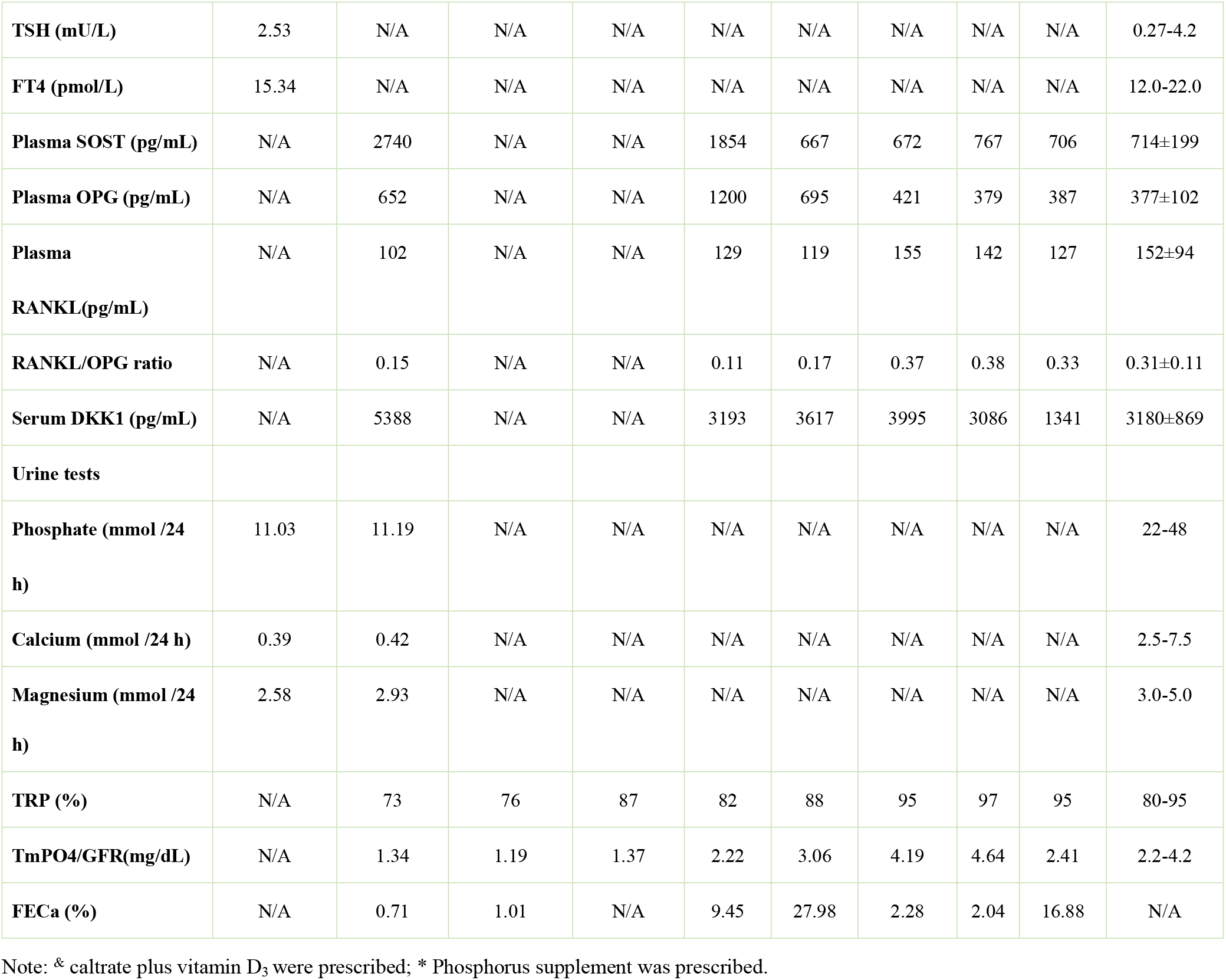
Clinical and Biochemical Results

### Circulating SOST, DKK1, receptor activator of RANKL and OPG levels

Plasma levels of SOST, RANKL and OPG were detected using kits from abcam (Cambridge, UK). Serum DKK1 levels were also measured using ELISA methods (R&D Systems, Inc., Minneapolis, USA). Twenty gender- and age-matched healthy adult males were selected as controls.

### Mutation analysis

WES of the proband was performed and the detected possible mutations were further analyzed in DNA samples from his parents, brother and children (KingMed Diagnostics, Guangzhou, China).

### Osteoclast culture

Osteoclasts from human peripheral blood were cultured as previously described ^10^. PBMCs were isolated using Ficoll-Hypaque Solution. Cells were washed in PBS twice, and plated on 24-well plates at a density of 1×106/well at 37°C in α-MEM, supplemented with 10% FBS, 1% penicillin/streptomycin and 25 ng/ml of macrophage colony stimulating factor (M-CSF) (R&D Systems, Inc., Minneapolis, USA). 6 days later, OC differentiation was induced with the medium supplemented with both 25ng/ml of M-CSF and 30ng/ml RANKL (R&D Systems, Inc., Minneapolis, USA). 7 days later, TRAP staining was performed using a kit from Sigma-Aldrich (sigma Chemical Co., St. Louis, MO, USA). TRAP-positive cells containing 3 or more nuclei were counted as OCs. 6mm*6mm bovine cortical bone slices were put into cell culture wells at the beginning of OC differentiation. 7 days after co-cultures, the slices were removed and evaluated for OC morphology and pit formation by scanning electron microscope (INCA PENTAFET X3, Oxford Instruments, Abingdon, Oxfordshire, UK).

### Plasmids

WT XLas cDNA was synthetized according to the sequence from Genebank database, and the mutant XLas cDNA was generated by site-directed mutant PCR and was confirmed by sequencing. The WT and mutant XLas cDNAs, tagged with enhanced green fluorescent protein (EGFP), were cloned into expression vector GV144 (Shanghai Genechem Co., Ltd., Shanghai, China).

### Expression and location of WT and Mutant XLas in SaOS2 cells

Human SaOS2 cells were maintained in αMEM containing 10 % FBS, 10 mM HEPES, 0.2 M L-Glutamine and penicillin/gentamycin at 37 °C with 5 % CO_2_. After reaching 70–80% confluence, cells were transfected with 1.5ug DNA per well with 1:3 ratio of Xtreme^HP^ transfection reagent (Roche Diagnostics Ltd., Indianapolis, USA). 72h after transfection, cells were fixed in 4% PBS-buffered paraformaldehyde at room temperature for 5min. Cell nuclei were dyed with DAPI.

72h after transfection, the growth medium was aspirated and the total RNA was extracted using a Trizol reagent (Thermo Fisher Scientific Inc., Waltham, USA). Relative gene expression levels were normalized to GAPDH, and analyzed with 2^-ΔΔCt^ method.

### cAMP production stimulated by PTH

72h after transfection, the confluent monolayer SaOS2 cells were serum starved for 6 hours prior to treatment. Cells were first treated with 3-isobutyl-1-methylxanthine (IBMX, 1mM) (R&D Systems, Inc., Minneapolis, USA) for 15min and then treated with PTH(1-34) (50 nM) (R&D Systems, Inc., Minneapolis, USA) for 20 min. Cells were lysed using 0.1M HCL and cAMP levels were measured using a direct cAMP ELISA kit (Sigma Chemical Co., St. Louis, MO, USA).

## Results

### Clinical and laboratory findings

The proband was prescribed caltrate plus vitamin D_3_ to exclude the secondary hyperparathyroidism induced by vitamin D deficiency. The serum levels of 25-hydroxyvitamin D_3_ were reversed within the normal range, however, elevated PTH and hypophosphatemia persisted and his serum calcium levels remained at the lower limit of the normal range. It could be seen from table 1 that the levels of B-ALP, OCN and CTX showed a tendency to decrease during the past 4 years. He had low TRP (%) (percent tubular reabsorption of phosphate) and TmPO4/GFR (tubular maximum phosphate reabsorption per glomerular filtration rate) (table 1), indicating reduced tubular reabsorption of phosphorus. Compared with the other members of his family, the proband showed lower Fractional Excretion of Calcium (FECa) (table 1), indicating low urinary calcium excretion. The DXA scan performed in February 2017 and October 2018 revealed progression in the bone density of the proband (table 1). X-ray of skull showed unclosed coronal suture and sagittal suture. Diffuse increase in bone mass was seen in the vertebral body. Pelvic X-ray showed bilateral femoral neck fractures and uneven bone density in pelvic. Cortical bone thickening was seen in long bones (Fig. 1A-D). His father had a history of fracture and was unable to walk for 5 years for unknown reasons. The proband or his family members did not show heterotopic ossifications in their skeletons.

**Figure 1.**
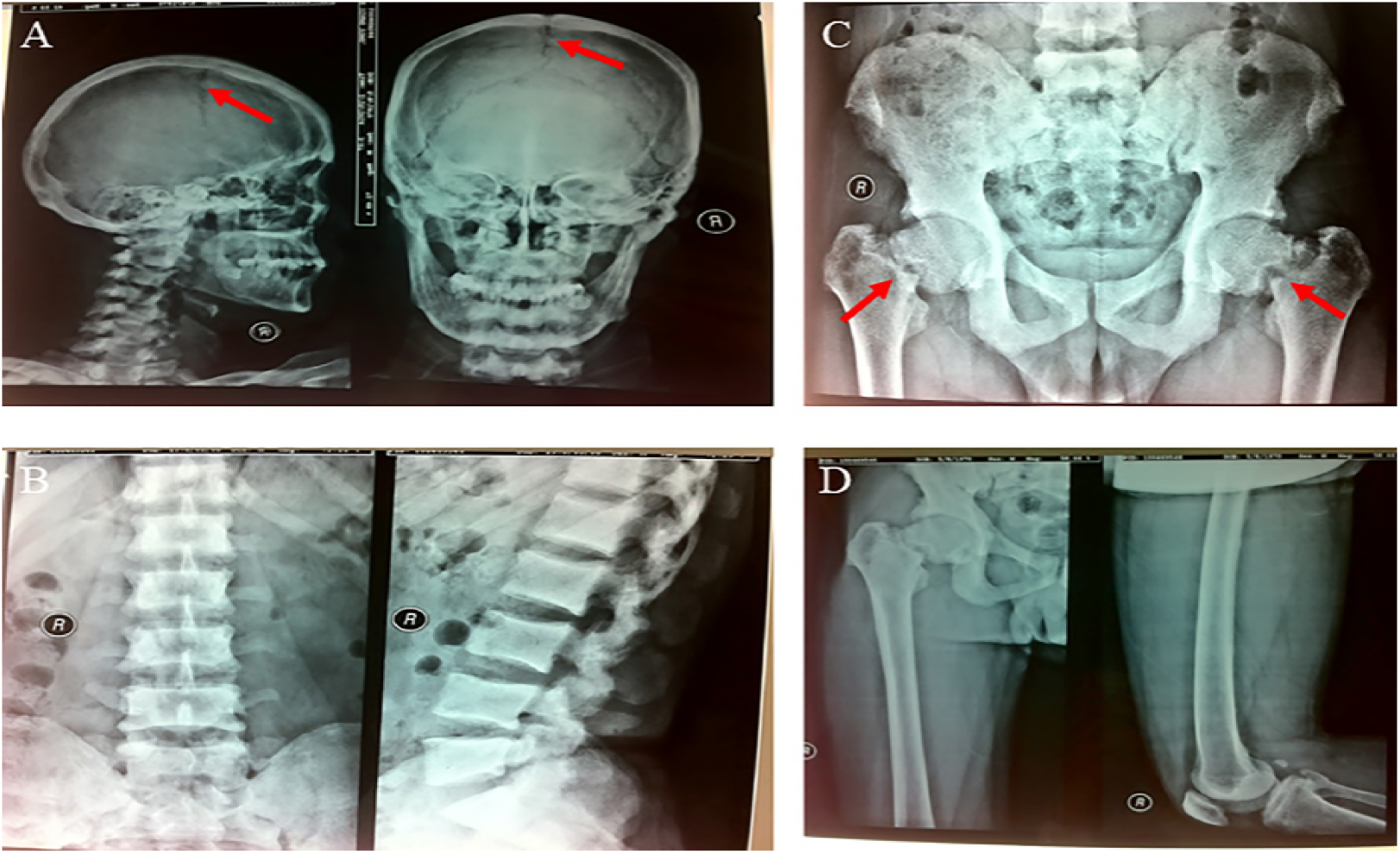
Radiographic features of this patient. (A). X-ray of skull showed unclosed coronal suture and sagittal suture (arrow). (B). Vertebral body showed diffuse increase in bone density. (C). Bilateral femoral neck fractures (arrow) and uneven density of pelvis. (D). Increased bone mineral density in femoral cortex.

Compared with the normal controls, this subject showed significantly higher SOST, OPG, and DKK1 levels, while lower RANKL levels and decreased RANKL/OPG ratio (table 1). His father also had higher SOST and OPG levels, as well as lower RANKL levels and decreased RANKL/OPG ratio (table 1).

### Mutational Analysis

Whole exome sequencing (WES) of the proband was performed. A novel heterozygous missense mutation c.424G>T (p. G142X) was found in exon 1 of GNAS isoform XLas (NM_080425.3). This changed the codon (GGA) for 142^nd^ amino acid glycine (G) to a stop codon (UGA) (Fig. 2A) and was predicted to be deleterious, leading to early termination of protein translation.This mutation was not found in OMIM, HGMD and Clinvar database, and was also not included in population databases, including 1000 Genomes, dbSNP, Exome Variant Server and ExAC Browser. This mutation was also found in his father and daughter. His mother, brother and son were negative.

**Figure 2.**
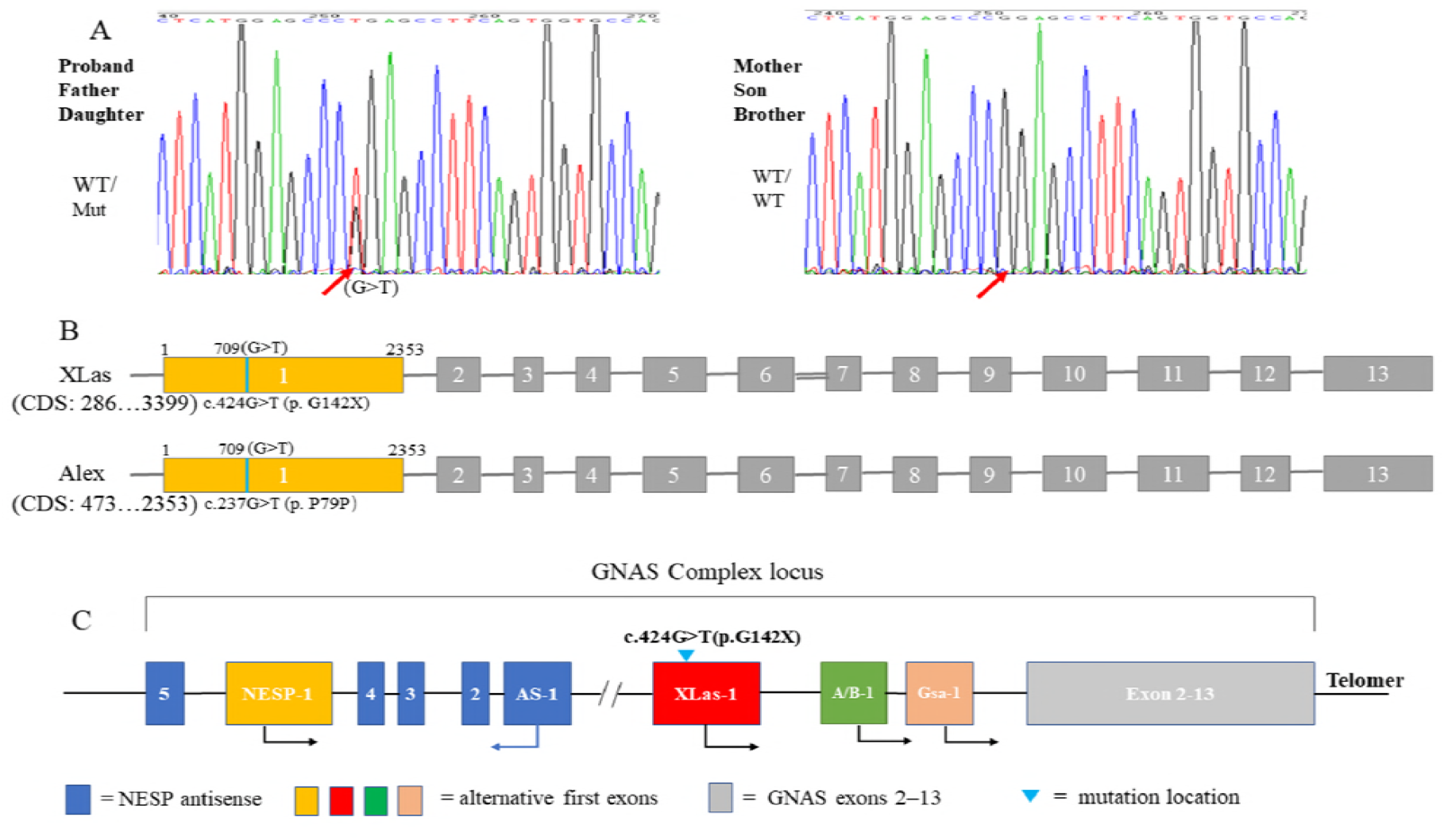
Genetic analysis of GNAS mutation. (A). A missense mutation c.424G>T (p. G142X) in XLas was found in proband, his father and daughter. (B). This mutation exclusively affects XLas, but not Gsα.

XLas, a long Gsα variant, uses an alternative first exon, which splices into exons 2–13 of Gsα. Alex (NM_001309883.1) is also generated from an alternative reading frame of the XLas transcript[9], which has no similarity to other proteins encoded by this gene. This mutation locates in the first exon of XLas, which is also located in the coding region of Alex and generates a synonymous mutation c.237G>T (p. P79P) (Fig. 2B). Gsα, NESP55 and A/B transcripts were predicted to be unaffected.

### Osteoclast(OC) formation and function

Osteoclasts were induced from human peripheral blood mononuclear cells (PBMCs) to oberserve the ability of in vitro osteoclast formation and bone resorption. TRAP staining reveal decreased osteoclast numbers induced from the PBMCs of the patient (Fig. 3G). However, scanning electron microscopy observation of the co-cultured bone slices showed normal osteoclast morphology and pit formation of the osteoclasts from the patient (Fig. 3C-F).

**Figure 3.**
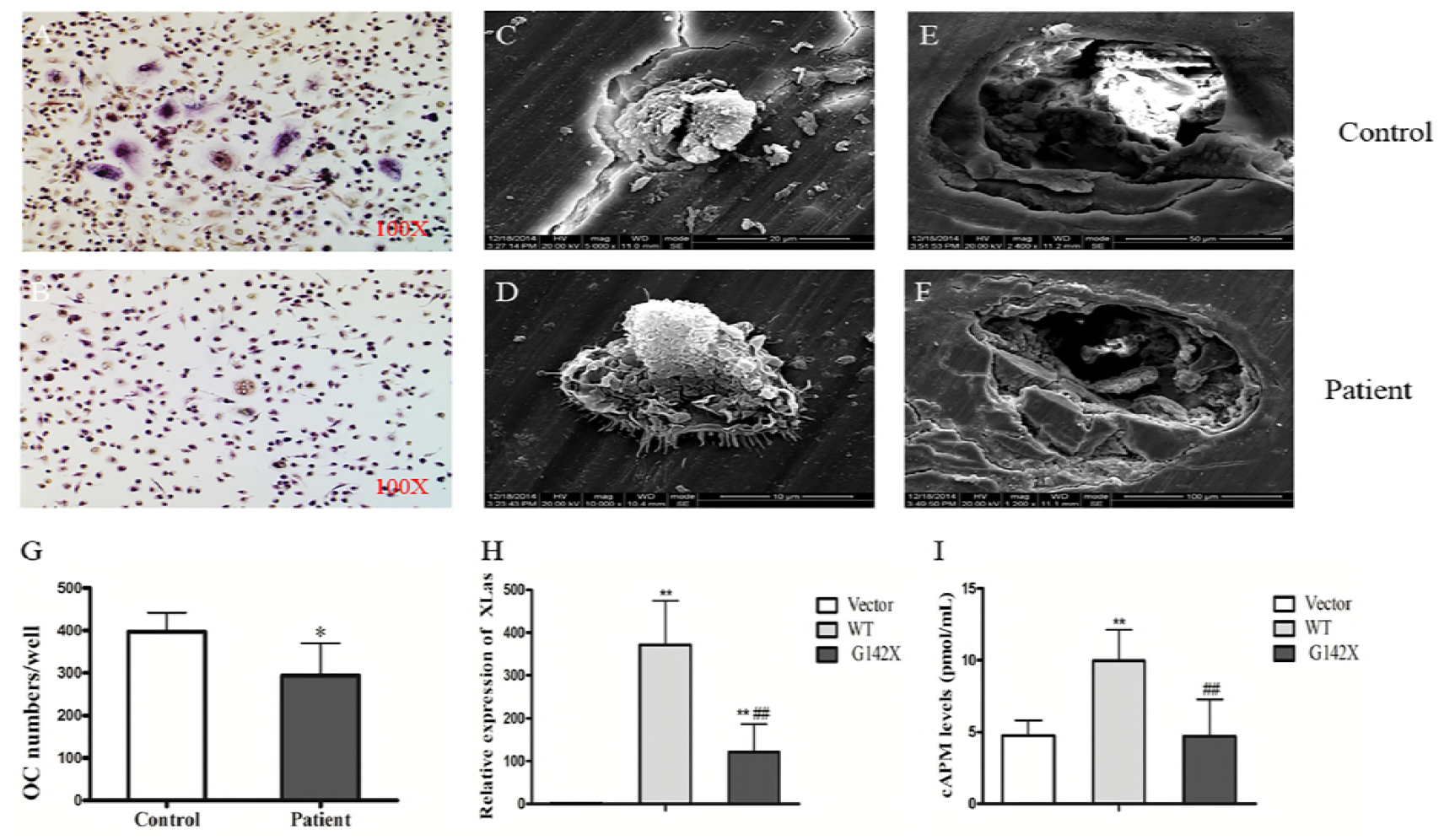
*In vitro* osteoclast culture and expression of mutant or WT XLas in SaOS2 Cells. (A-B). TRAP staining of osteoclasts. (C-D). Adherent cells on the bone slices. (E-F) Pit formation after ultrasonic removal of the adherent cells. (G). The number of osteoclasts in the patient culture was significantly lower than that in the control culture. **p<0.01, compared with control. (H). The mRNA expression of WT and mutant XLas was significantly increased in the transfected SaOS2 cells. **p<0.01, compared with vector; ## p<0.01, compared with WT. (I). The production of cAMP induced by PTH in SaOS2 cells transfected with WT XLas was significantly higher than that in the empty vector group, which was blunted in G142X mutation. **p<0.01, compared with vector; ## p<0.01, compared with WT.

### Expression and location of Wild type (WT) and mutant XLas in SaOS2 cells

WT and mutant XLas cDNA were transfected into human SaOS2 cells. In these transfected cells, EGFP-tagged WT XLas was localized to the cellural membrane and cytoplasm. On the contrary, the expression of EGFP-tagged mutant XLas was absent in transfected SaOS2 cells (Fig. 4). The mRNA expression of WT and mutant XLas was significantly higher in transfected cells than in cells transfected with empty vector (Fig. 3H).

**Figure 4.**
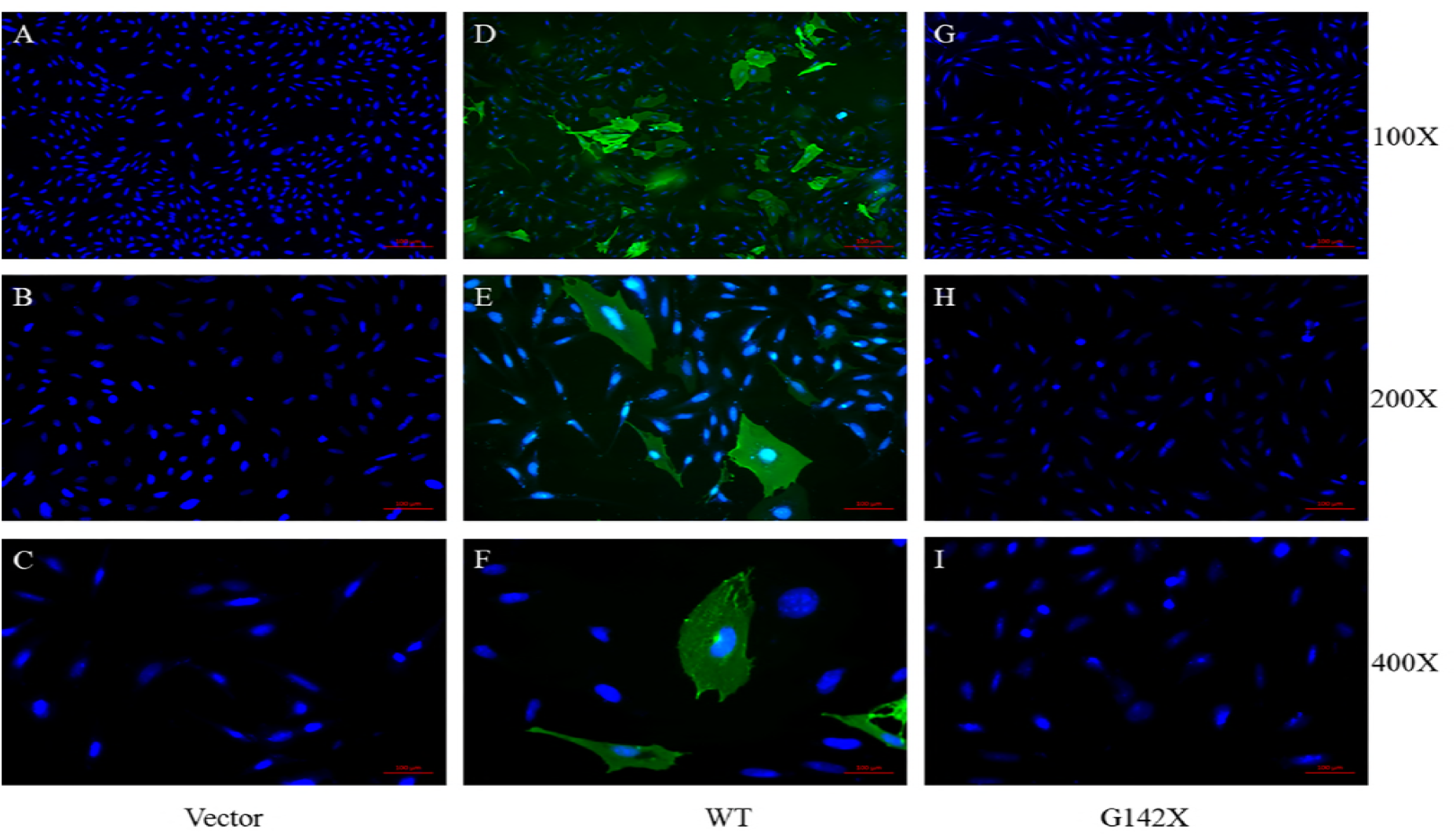
(A-C). SaOS2 cells transfected with empty vector. (D-F). SaOS2 cells transfected with WT XLas. EGFP-tagged WT XLas was localized to the plasma membrane and cytoplasm. (G-I). SaOS2 cells transfected with mutant XLas.

### cAMP production

The ability of cAMP generation under PTH stimulation in transfected cells were determinded. SaOS2 cells transfected with WT XLas showed higher cAMP production after PTH stimulation, while SaOS2 cells transfected with mutant XLas showed similar lower cAMP production as that in SaOS2 cells transfected with empty vector (Fig. 3I).

## Discussion

This is the first report of a mutation located in the first exon of XLas in humans, which was predicted to have no significant effects on other gene products of this locus. This novel mutation causes a new constellation of clinical features including high bone mass, unclosed cranial suture, fractures, hypophosphatemia, and elevated PTH levels. By contrary, the major clinical features of AHO, caused by heterozygous inactivating mutations in Gsα-coding GNAS exons, include obesity with round face, short stature, brachydactyly, subcutaneous ossification, and mental retardation [4]. In some cases, TSH, gonadotropins and GHRH resistance may be variably present [10]. Previously reported mutations often disrupt multiple protein-coding transcripts in addition to that encoding Gsα [11, 12], making it difficult to distinguish the contributions of each transcript to disease phenotypes. Therefore, this case is very rare and precious. Our study indicates XLas plays a key regulatory role in bone metabolism and may be involved in PTH/cAMP signaling pathway in physiological conditions.

Gsα is able to stimulate the production of the second messenger cAMP under PTH stimulation. However, the significance of XLas still remains largely unknown in humans with conflicting results [6, 8, 13]. The abnormal skeletal phenotype in this patient implies that XLas has an important physiological role in humans. Our *in vitro* study clearly demonstrated that WT XLas was capable of stimulating cAMP production in human SaOS2 cells, which was blunted by G142X mutation. PTH shows both anabolic and catabolic actions on the skeleton, depending on the ways in which PTH is administrated [14]. Through targeting cAMP/PKA pathways, PTH favors bone resorption by stimulating RANKL while inhibiting OPG expression in osteoblasts/osteocytes via the PTH1R, rendering increased RANKL/OPG ratio [15–17]. Also through cAMP/PKA signaling pathway, PTH induces bone formation, at least in part, by its ability to downregulate SOST and DKK1 expression in osteocytes [14, 18–21]. This proband showed increased circulating leves of SOST, DKK1 and OPG, while decreased RANKL levels and reduced RANKL/OPG ratio, concurrently with the elevated PTH levels, indicating impaired skeletal response, at least partly, to PTH as a result of lower cAMP production induced by G142X mutation (Figure 5). Our study suggested that XLas may exert similar regulatory effects as Gsα regarding the expression of target genes as well as cAMP production. There is another possibility that the observed phenotype may be primarily resulted from XLαs deficiency in bone but not from the action of PTH. However, based on our *in vivo* and *in vitro* studies, especially the changes of circulating SOST, DKK1, RANKL and OPG levels, it is more likely to be secondary to the impaired PTH/cAMP pathway in the skeletal tissue.

**Figure 5.**
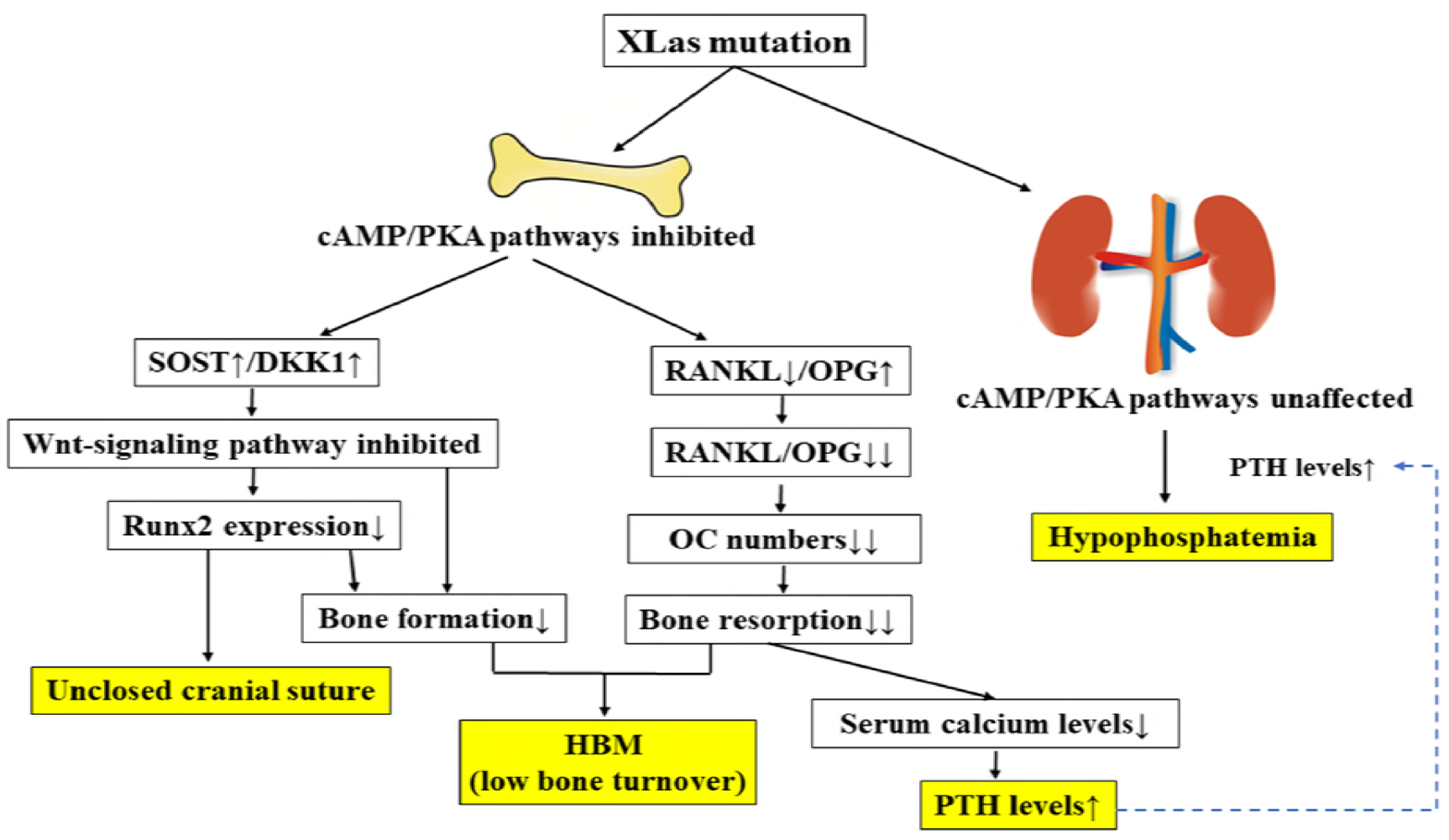
XLas mutation leads to selective skeletal resistance but normal renal tubular response to PTH.

Although SOST, DKK1, RANKL and OPG have not been established as the clinical bone markers in humans, previous studies have suggested that disorders of parathyroid influence the circulating levels of these factors. Serum SOST levels are decreased in primary hyperparathyroidism and increased in hypoparathyroidism [22, 23], while serum DKK1 levels are increased in hyperparathyroidism [24]. Elevated RANKL/OPG ratio has also been found in subjects with hyperparathyroidism [25, 26]. However, no related studies have been reported in subjects with PHP. Except for serum DKK1 levels, the elevated plasma SOST levels and reduced RANKL/OPG ratio may indicate decreased parathyroid function in proband and his father.

This patient showed a progressive increase in BMD, and the underlying mechanism was speculated to be due to impaired bone resorption. First, *in vitro* study found decreased osteoclast numbers induced from the PBMCs of this patient. Second, circulating levels of RANKL and the RANKL/OPG ratio were significantly reduced in patient, which may lead to lower osteoclast formation. Furthermore, osteopetrosis cases due to impaired bone resorption tend to have increased fracture risks, while sclerosteosis cases secondary to enhanced bone formation do not [27]. This patient showed bilateral femoral neck fractures, further confirming the impaired bone resorption as a result of reduced osteoclast formation may be the main casue of HBM.

The most perplexing thing about this patient is his bone turnover status. There are some contradictions in his clinical manifestations, bone metabolism index and the circulating levels of SOST, DKK1, RANKL and OPG. Bone biopsy remains the gold standard to assess the true bone metabolism status. However, bone biopsy was not performed due to the poor bone quality of this patient. It is well known that primary hyperparathyroidism causes increased bone turnover and reduced BMD, especially at the cortical bone [28]. On the contrary, idiopathic hypoparathyroidism (IHP) cases often have reduced bone turnover and increased BMD than the general population [29, 30]. Compared with IHP and nonsurgical hypoparathyroidism (Ns-HypoPT) patients, PHP subjects, especially PHP type 1B, tend to have lower BMD, indicating incomplete skeletal resistance to PTH may exist [31–33]. However, when compared with normal controls, patients with PHP1a showed a significantly greater total body BMD [34]. Furthermore, striking osteosclerosis has been found in two brothers diagnosed with PHP1b [35]. Gsa in skeleton is biallelically expressed. Therefore, no matter the origin of mutation is, GNAS mutation may lead to decreased skeletal response to PTH [36, 37]. It is reasonable to infer that bone turnover and bone metabolism status in PHP are closer to hypoparathyroidism than to hyperparathyroidism. However, the levels of bone metabolism indicators were significantly high in this patient. In other two studies, the levels of bone turnover markers were also high in subjects with PHP [31, 33]. It is notable that the proband’s bone indicators, including B-ALP, OCN and CTX, showed a tendency to decrease during the past 4 years, despite the progressive increase in BMD. His persistent hypophosphatemia may play a role in the elevated B-ALP levels.

Our *in vitro* study clearly showed reduced osteoclast formation ability in this patient, which is consistent with the reduced RANKL/OPG ratio. Based on our experience, *in vitro* osteoclast culture could reliably reflect the osteoclast formation ability of humans [38]. Although this patient had increased CTX levels, we believe he had inhibited bone resorption as a result of decreased osteoclast formation, which is also consistent with osteopetrosis. As for bone formation, this patient had increased levels of SOST and DKK1, both of which could inhibit Wnt-mediated bone formation [27]. Furthermore, bone formation is coupled to bone resorption in general. Therefore, it is reasable to infer that both bone resorption and bone formation were inhibited in this patient. The main physiological function of PTH is to promote bone resorption [5]. The impaired skeletal response to PTH may result in more inhibition on bone resorption than on bone formation, leading to increased bone mass (Figure 5). Therefore, we believe this patient belongs to a type of HBM with low bone turnover, primarily resulting from impaired bone resorption. Our study also suggests that exogenous RANKL may alleviate the progressive increase in bone mass in this patient.

In the classic form of PHP, the PTH resistance is confined to the renal proximal tubule, leading to hypocalcemia, hyperphosphatemia, and elevated levels of PTH levels [39]. In fact, PTH resistance can occur either in the renal tubular level or the skeleton [39]. Given that the proband showed elevated serum PTH concurrently with increased phosphate excretion, it appears that PTH resistance is not present in the proximal tubule. In addition, urinary calcium excretion was extremely low, consistent with elevated PTH and a lack of resistance in distal nephron. In renal proximal tubules, Gsα is expressed only from the maternal allele, which may also apply to XLas based on patient-specific clinical data. Therefore, in this proband, the renal tubular response to PTH may be normal. We speculated the paternally inherited mutation located in XLas impaired skeletal response to PTH, leading to reduced serum calcium and elevated PTH level. The persistently elevated PTH caused hypophosphatemia. As a compensation, his serum calcium levels were maintained at the lower limit of the normal range.

It is notable that the patient showed unclosed coronal suture and sagittal suture. Unclosed cranial suture has also been reported in patients carrying mutations in runt-related transcription factor 2 (Runx2), which is known to affect metopic suture fusion and plays a key role in osteoblast differentiation mediated by Wnt-signaling pathway [40]. Runx2 mutations have been described in subjects with cleidocranial dysplasia (CCD) (MIM 119600), which is a rare hereditary skeletal disorder [40, 41]. WES has excluded Runx2 mutation in this patient. However, our study found increased circulating levels of SOST and DKK1 in this patient, both of which could inhibit Wnt-pathway mediated bone formation. Runx2 is one of the major target genes of activated Wnt-pathway [42], and the inhibited Wnt-signaling pathway may lead to reduced Runx2 expression. Therefore, it is reasonable to speculate that the unclosed cranial suture may be secondary to the impaired Wnt signaling and the resultant inhibited Runx2 expression (Figure 5).

His father also carried this mutation and the origin of mutant allele was unknown. It seemed that his father manifested a similar but milder phenotype. His daughter also carried the same mutation and presently showed no similar synptoms, possibly due to her young age or incomplete penetrance. It is suggested that XLas may be essential for normal fetal growth and development, and paternal mutations in XLas may lead to severe intrauterine growth retardation [11]. The birth records of this patient and his father were unknown. The daughter was born at term with a birth weight of about 2.6 kg and was well after birth. This family will be regularly followed up.

In conclusion, we first report a novel mutation located in the first exon of XLas in a patient with HBM, unclosed cranial suture, hypophosphatemia, and elevated PTH, indicating there is still a lot of ignorance about the physiopathologic roles of altenative GNAS gene products. The identification of cases with novel genetic and epigenetic defects indicates the urgent need for a new classification of this spectrum of diseases [5]. Our findings further expand the spectrum of clinical manifestation of diseases due to GNAS mutaions, and also urge further investigation to explore the regulatory role of XLas in bone metabolism. Our study indicates that GNAS locus should be considered as a candidate gene for HBM.

## Acknowledgments

This work was supported by grants from the National Natural Science Foundation of China (No. 81770875, 81572639, 81370969 to Xijie Yu) and the Science & Technology Department of Sichuan Province (2018SZ0142 to Xijie Yu). This study was also supported by the National Natural Science Foundation of China (grant no. 81702156 to Y Meng), Postdoctoral Science Foundation of China (grant no. 2017M61060 to Y Meng) and Postdoctoral Research Foundation of Sichuan University (grant no. 2017SCU12038 to Y Meng). The authors thank Miss Xiao Yu from the University of Michigan, for her help with editing the language.

## Authors’ roles

Study design: Xiang Chen and Xijie Yu. Study conduct: Xiang Chen, Ying Xie, Shan Wan, Li Li, Jie Zhang and Bo Su. Data collection: Xiang Chen and Yang Meng. Data analysis: Xiang Chen. Data interpretation: Xiang Chen and Xijie Yu. Drafting manuscript: Xiang Chen. Revising manuscript content: Xijie Yu. Approving final version of manuscript: Xiang Chen and Xijie Yu. Xiang Chen and Xijie Yu take responsibility for the integrity of the data analysis.

